# Revisiting G3BP-S149 phosphorylation and its impact on stress granule assembly

**DOI:** 10.1101/598201

**Authors:** Marc D. Panas, Nancy Kedersha, Tim Schulte, Rui M. Branca, Pavel Ivanov, Paul Anderson

**Affiliations:** Harvard Medical School and Brigham and Women’s Hospital; Division of Rheumatology, Immunology and Allergy; 60 Fenwood Road, Boston, MA 02115; Karolinska Institutet; Department of Microbiology, Tumor and Cell Biology; Nobels väg 16, Stockholm SE-171 77, Sweden; Science for Life Laboratory, Department of Medicine Solna, Karolinska Institutet, Stockholm SE-171 77, Sweden; Clinical Proteomics Mass Spectrometry, Department of Oncology-Pathology, Science For Life Laboratory and Karolinska Institutet, Stockholm, Sweden

## Abstract

Stress granules (SGs) are cytoplasmic, non-membranous RNA/protein structures that assemble in response to environmental stress. G3BP is a critical SG-nucleating protein, and its ability to regulate SGs has been reported to be regulated by serine 149 phosphorylation. We now report that the constructs engineered to contain non-phosphorylatable and phosphomimetic (G3BP1-S149A and G3BP1-S149E, respectively) mutations used in many studies include additional unintended mutations (A54T/S149A and S99P/S149E) one of which (S99P) is responsible for the effects on SG assembly attributed to S149E. Specifically, the S99P mutation alone reduces SG nucleation and impairs the ability to rescue SG assembly in ΔΔG3BP1/2 U2OS KO cells, challenging the widely-stated conclusion that de-phosphorylation of serine 149 in G3BP1 promotes SG assembly. We used comparative mass spectrometry analysis of both (1) ectopically expressed GFP-G3BP1 in ΔΔG3BP1/2 U2OS KO and (2) endogenous G3BP1 in wild-type U2OS, with and without sodium arsenite treatment, in an attempt to reproduce earlier findings, but found no significant changes in S149 phosphorylation that correlate with arsenite-induced SG formation.

## Introduction

Sudden changes in environmental conditions, such as heat shock, oxidative stress, UV or viral infection, affect all cells, and can damage proteins, lipids and nucleic acids (Anderson and Kedersha, 2008; Buchan and Parker, 2009). Cells require protective programs to maintain viability in the face of changing environmental conditions, in part by reprogramming protein expression (Buchan and Parker, 2009), and by reducing expression of housekeeping proteins to enhance production of pro-survival proteins (Kedersha and Anderson, 2002). Stressed cells inhibit translation initiation, thus increasing the amount of stalled 48S preinitiation complexes (PICs) (Panas et al., 2016) released from polysomes. These PICs condense to form non-membrane-enclosed foci known as stress granules (SGs) (Kedersha et al., 2000). Core components of SGs are 48S PICs and mRNA, but they also enriched for a large number of RNA-binding proteins that promote SG condensation, such as TIA1/R, FMRP/FXR1, Caprin1 and G3BP. In addition to these RNA-binding proteins, an eclectic collection of signaling molecules are recruited into SGs, including RACK1, PKCα, USP10, and TRAF2 (Kedersha et al., 2013).

SGs are dynamic structures into which protein contents rapidly shuttle in and out, although SGs themselves can persist for hours (Kedersha et al., 2000). This property and the lack of a limiting membrane have led to the proposal that SG formation is mediated by a liquid/liquid phase transition (Han et al., 2012; Kato et al., 2012; Weber and Brangwynne, 2012) by which stalled PICs are condensed into visible subcellular regions by the activity of protein aggregases (Kedersha and Anderson, 2007; Kedersha et al., 2016; Reineke et al., 2015). The threshold for SG formation is determined by two general events: the bulk amount of stalled PICs released from polysomes, and the local concentration/aggregation of SG-nucleating proteins. Stalled PIC levels are regulated by stress-activated <u>e</u>ukaryotic <u>i</u>nitiation <u>f</u>actor (eIF)2α kinases, reducing available ternary complexes and resulting in stalled 48S PICs (Kedersha et al., 2002; Kedersha et al., 1999). Polysome disassembly resulting in SG formation is also caused by drugs such as pateamine A (Bordeleau et al., 2006a; Dang et al., 2006) or hippuristanol (Bordeleau et al., 2006b; Tsumuraya et al., 2011), compounds that target the eIF4F complex, which leads to stalled 48S PICs and thus triggers SG formation.

Ectopic expression of SG-nucleating proteins, such as TIA1 or G3BP, induces the formation of SGs even in the absence of drugs or stress (Kedersha and Anderson, 2007) by increasing the rate at which available PICs are condensed into SGs. SG-nucleating proteins are rich in prion-like, low complexity (LC), and unstructured/intrinsically disordered (ID) protein regions (Uversky, 2017) that mediate protein aggregation (Tompa, 2005). The SG protein TIA1 contains a prion-like domain that can form insoluble aggregates (Gilks et al., 2004). The SG-nucleators G3BP1 and G3BP2 (hereafter referred to jointly as G3BP) lack a prion-like domain but contain extensive disordered regions, adjacent to an ordered NTF2-like dimerization domain (Tourriere et al., 2003) that is essential for SG assembly. The SG-inducing activity of G3BP is inhibited by interactions with other proteins such as USP10, which contains a short motif (FGDF) that binds the G3BP NTF2-like domain. The FGDF motif is also found in the viral non-structural protein nsP3 of Semliki Forest virus that binds G3BP (Panas et al., 2015; Schulte et al., 2016). Overexpression of these FGDF-containing proteins blocks SG formation (Kedersha et al., 2016; Panas et al., 2015).

The NTF2-like domain mediates dimerization of G3BP1, which is reportedly regulated by phosphorylation of a site (Ser 149) located in a disordered acidic region adjacent to the NTF2 domain (Tourriere et al., 2003). Serine 149 was described to be constitutively phosphorylated, and de-phosphorylated in response to sodium arsenite (SA) treatment, concurrent with SG formation. The authors proposed that constitutive phosphorylation of S149 prevents G3BP1 dimerization and consequently inhibits SG formation; its stress-induced de-phosphorylation would then allow dimerization and promote SG formation. Evidence to support this model was obtained by using overexpressed non-phosphorylatable G3BP1 mutant S149A and phosphomimetic mutant S149E, which showed that S149A G3BP1 efficiently nucleated SGs while S149E failed to do so (Tourriere et al., 2003). A subsequent study from our lab using stably expressed S149A and S149E G3BP1 mutants in ΔΔG3BP1/2 double knockout U2OS cells indicated that the S149E mutant is impaired in SG assembly (Kedersha et al., 2016).

Here we report that there are additional mutations in the original pEGFP-C1-G3BP1-S149A and S149E constructs (Tourriere et al., 2003) that were generously shared and widely disseminated. These constructs were used in our study (Kedersha et al., 2016) and many other studies (Barr et al., 2013; Jedrusik-Bode et al., 2013; Kedersha et al., 2016; Kwon et al., 2007; Sahoo et al., 2018; Sahoo et al., 2012; Szaflarski et al., 2016). We have now identified second mutations within the NTF2-like domains of the original constructs. Specifically, we identified an alanine to threonine conversion at position 54 in the S149A (actually A54T/S149A) construct, and a serine to proline conversion at position 99 in the S149E construct (actually S99P/S149E). Whereas the single A54T mutation has no effect on SG assembly, the single S99P mutation destabilizes the protein, dominantly inhibits SG formation, and only partly rescues SG formation in G3BP1/2 KO cells. Transient transfections and immunoprecipitations reveal that G3BP1-S99P does interact with USP10 and Caprin1, but exhibits impaired dimerization. It is less efficiently expressed owing its ubiquitination and instability, and co-aggregates with the autophagy adaptor protein sequestosome 1 *in vivo*. Mapping of the S99P mutations to available crystal structures supports the observed impairment of dimerization. Mass spectrometry (MS) analysis of S149 phosphorylation in untreated versus SA treated cells reveals no significant differences, indicating that S149 phosphorylation does not correlate with SG formation. Finally, repaired S149A and S149E constructs lacking these additional mutations behave indistinguishably from wt G3BP1 in their ability to mediate SG assembly.

These data show that the impaired ability of the original pEGFP-C1-G3BP1-S149E construct to nucleate or rescue SGs is caused by the accidental S99P mutation, rather than the intentional S149E mutation. This finding impacts other reports that used these plasmids to investigate the functional effects of S149 phosphorylation of G3BP1.

## Results

### G3BP1-S149E contains a second S99P mutation localized within NTF2-like domain of G3BP1

We set out to examine how S149 in the acidic region could regulate presumed NTF2-like domain dimer formation, using a 1-168 fragment of G3BP1 previously described (Kedersha et al., 2016). After re-cloning G3BP1-168-wt, G3BP1-168-S149A and G3BP1-168-S149E into bacterial expression vectors, we encountered unexpected issues when purifying His-tagged G3BP1-168-S149E (Fig. S1A). While both wild-type and the S149A mutant migrated as soluble proteins with defined peaks in size exclusion chromatography (SEC), the S149E protein appeared as a major peak in the void volume, but not at the expected molecular weight (Fig. S1A). Therefore, we checked the sequences of the bacterial expression plasmids and discovered secondary mutations of Ala-54 to Thr (A54T) and Ser-99 to Pro (S99P) in the G3BP1-S149A and G3BP1-S149E mutants, respectively. We then identified these same mutations in the original stocks of pEGFP-C1-G3BP1-S149A and S149E (Tourriere et al., 2003) that we had obtained in 2004. We then requested and received a second batch of plasmids (in 2016), and confirmed that both sets of plasmids harbored the same secondary mutations. The mutant constructs are diagramed in Fig. 1A.

**Figure 1:**
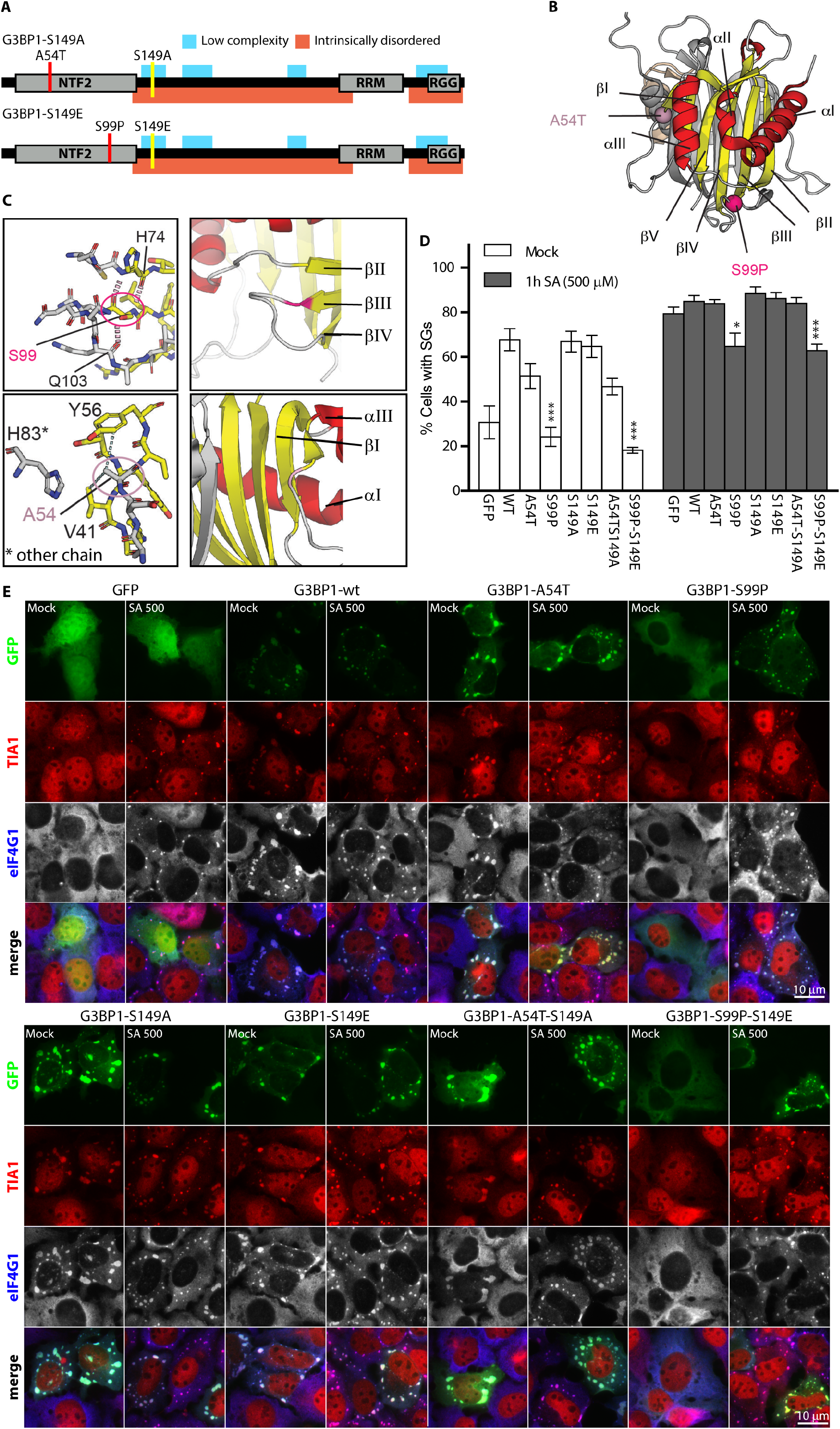
G3BP1-S99P and G3BP1-S99P-S149E mutants exhibit both impaired SG nucleation and recruitment to SGs in U2OS-wt cells. (A) Pictogram of GFP-G3BP1-S149A showing the additional mutation A54 to Thr (red) or GFP-G3BP1-S149E showing the additional mutation at position S99 to Pro (red). The S149 site is indicated in yellow. Gray boxes represent structured classical domains, and red areas indicate ≥50% predicted ID/IU regions, whereas aqua shading represent LC-regions. (B) Crystal structure of the NTF2-like domain of G3BP1. The Cα atoms of the two accidental secondary mutations A54T and S99P are highlighted as spheres colored in light pink and pink, respectively. Strands, helices and loop regions are colored in yellow, red and light grey. The second unit of the dimer is shown in grey. (C) Close-up view of the mutated residues. Peptides are represented in stick format, oxygen and amino-groups in blue and red, respectively. Carbons are colored according to the color code for strands, helices and loops as in (B). Hydrogen bonds and hydrophobic contacts of the mutated residues are illustrated as red and grey dashes. The cartoon figures in the right panels are shown for better orientation with same color code as in (B). (D) Quantification of SG data in (E), using TIA1 and eIF4G1 as SG markers. Data shown are mean +/- SEM and analyzed using unpaired t-test. *, P < 0.05; ***, P < 0.005; n = 5. (E) Localization of GFP-tagged proteins (green) in transiently-transfected U2OS cells, untreated (Mock) or treated with 500 μM sodium arsenite (SA) for 1h prior to fixing and staining for endogenous TIA1 (red) and eIF4G1 (blue in merged view, grey in separate channel). Bar, 10 μm.

Structural analysis of the crystal structure reveals that Ala-54 is localized in a loop region preceding the αIII helix, in hydrophobic contact with Tyr-56 and Val-41. While its substitution to Thr most probably has only minor effects, its close proximity to His-83 of the neighboring chain could affect G3BP1 dimerization (Fig. 1C), possibly explaining the observed altered mobility of the A54T/S149E mutation relative to wt G3BP1 in size exclusion chromatography (Fig. S1A). In contrast, Ser-99 is localized at the end of strand βIII, embedded in a β-sheet-mediated hydrogen bond pattern with His-74 of strand βII, and its side chain hydroxyl-group is fixed by another hydrogen bond to the carbonyl backbone of Gln-103. Substitution of Ser-99 to Proline has major consequences, as it disrupts the hydrogen-bond pattern, and also rigidifies the peptide backbone (Fig. 1C). An *in silico* substitution of Ser-99 to proline renders the formerly favored Phi and Psi angles of the Ser-99 backbone (Fig. S1B, panel 1, green arrows) into disallowed regions of the proline backbone in Ramachandran plots (Fig. S1B, panel 4, red arrows). In addition, we used the RosettaBackrub server to predict residues that are tolerated at these positions, and to model structures of the A54T and S99P mutants (Davis et al., 2006; Davis et al., 2007; Lauck et al., 2010; Smith and Kortemme, 2011). While both wild-type residues are highly favored (A54 and S99) at their respective positions, Pro-99 is less favorable as compared to Thr-54 (Fig. S1C, red boxes). The modelled structures of the S99P mutant exhibit a significant conformational adjustment of more than 2.5Å of the associated loop region, while the modeled structures of the A54T mutant are almost identical in their backbone conformation adjustment of 0.2 Å as compared to the wild-type structure (Fig. S1D). Thus, we conclude that the S99P mutation was the main cause of our failed protein purification attempt for the presumed S149E variant. We then hypothesized that the same S99P mutation may account for *in vivo* phenotypes ascribed to S149E, as the “S149E” construct was really the double mutant S99P/S149E. To test this hypothesis, we ascertained the effects of each single mutation on SG formation, and also revisited the phosphorylation state of G3BP-S149 during stress using mass spectrometry (MS).

### S99P disrupts SG condensation while S149E does not

We transfected U2OS-wt cells with the indicated constructs and assessed SG formation with/without sodium arsenite (SA) treatment (Fig. 1D, 1E). We counterstained the transfected cells for SG markers TIA1 and eIF4G1, to confirm bona fide SG formation (Fig. 1E). Cells transfected with GFP-tagged G3BP1-wt, A54T, S149A, S149E, and A54T-S149A displayed comparable rates of SG formation, but cells transfected with S99P and S99P-S149E exhibited fewer SGs than the other forms of G3BP1 and fewer than the GFP only control (Fig. 1D, 1E), consistent with the results obtained in COS7 cells reported earlier (Tourriere et al., 2003). In cells treated with SA, all GFP-G3BP1 variants are recruited to SGs. G3BP1 variants containing the S99P mutation significantly repress both spontaneous and SA-induced SGs (Fig. 1D, E), in partial agreement with the original study, which showed that G3BP1-(S99P)-S149E inhibited spontaneous but not SA-induced SGs.

We next assessed the extent to which the various forms of G3BP1 can rescue SGs in ΔΔG3BP1/2 U2OS cells, which lack endogenous G3BP1 and G3BP2 proteins and are unable to form SGs in response to most stresses. SG competence is rescued by reconstituting these cells with exogenously-expressed G3BP1, either transiently or stably (Kedersha et al., 2016). Transient transfection of ΔΔG3BP1/2 U2OS cells reveals that G3BP1-wt, A54T, S149E and A54T-S149A all rescue the formation of SGs to comparable levels (Fig. 2A and B). A very modest increase in SG nucleation without stress was observed upon transfection with the S149A mutant (Fig. 2B, * columns). However, variants bearing the S99P mutation were significantly impaired in their ability to nucleate (without stress) or rescue (with stress) SG formation (Fig. 2A, quantified in 2B, ** columns). In addition to SA, other stresses (clotrimazole and pateamine A) gave similar results (Fig. S2). Constructs bearing the S99P mutation were significantly less effective at rescuing SGs under all stress conditions (Fig. 2B, ** columns).

**Figure 2:**
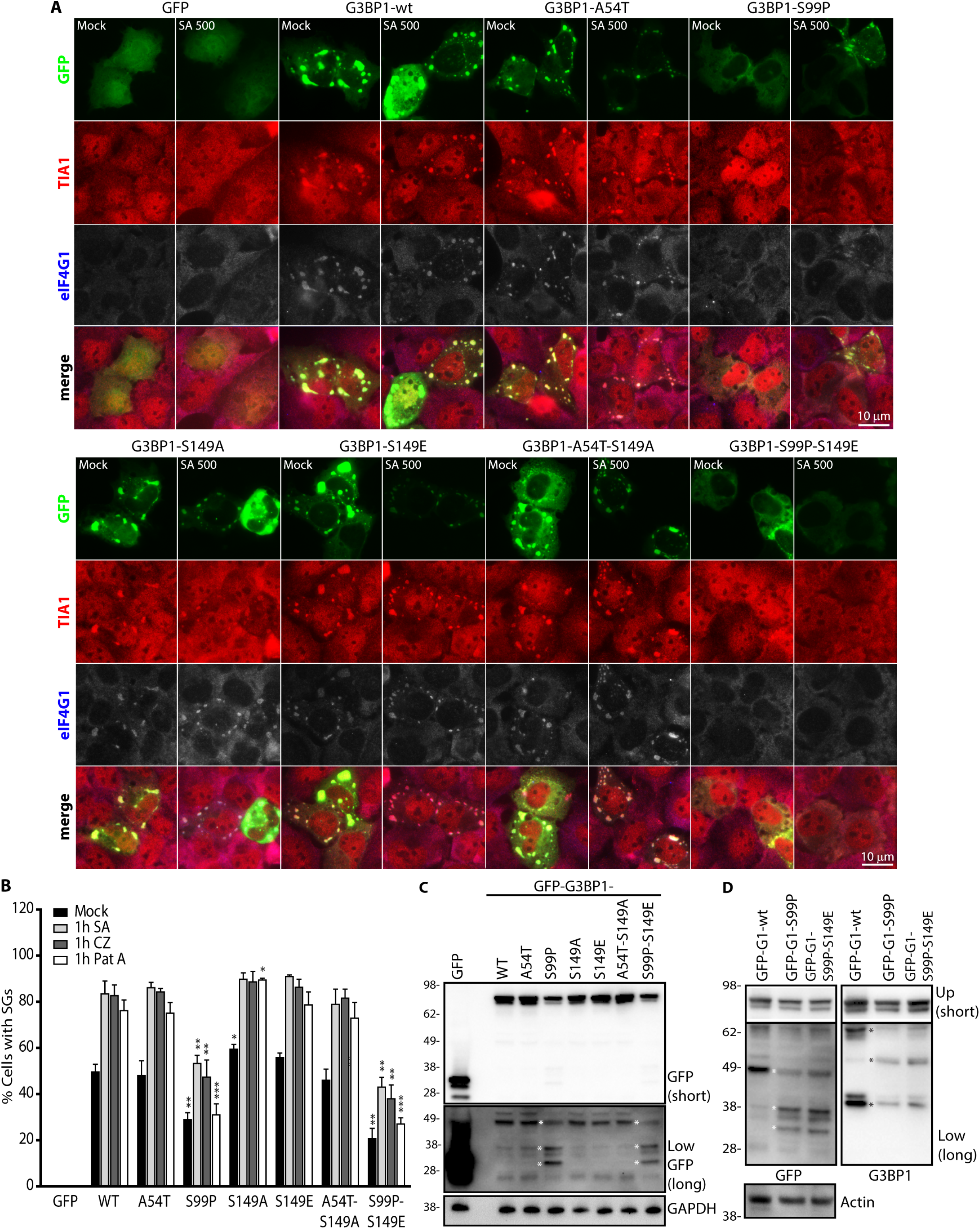
G3BP1-S99P-containing mutants exhibit reduced SG rescue in ΔΔG3BP1/2 cells. (A) Cells were transiently transfected with the indicated GFP-tagged (green) constructs, then untreated (Mock) or treated with 500 μM sodium arsenite (SA) for 1h. Cells were fixed and stained for endogenous TIA1 (red) and eIF4G1 (blue, represented as grey). Bar, 10 μm. (B) Quantification of SGs in cells shown in A, scored using TIA1 and eIF4G1 as SG markers. Data shown are mean +/- SEM and analyzed using unpaired t-test. *, P < 0.05; **, P < 0.01; ***, P < 0.005; n = 4. (C) Western blot of transfectants shown in (A). Lower portion, longer exposure. * indicates altered GFP products. (D) Transiently overexpressed constructs in ΔΔG3BP1/2 U2OS cells, blotted as indicated. Upper (Up) membrane exposure shorter than the lower part (Low) of the membrane. * indicates altered GFP or G3BP1 products in the mutants.

### Increased proteolysis of G3BP1-S99P impairs SG formation

The reduced ability of S99P constructs to rescue SGs could be due to lower expression levels. To assess this, we transiently transfected G3BP1 constructs into ΔΔG3BP1/2 U2OS and quantified the protein levels using western blotting. Consistently, the constructs harboring the S99P mutation showed reduced protein expression relative to G3BP1-wt and the other mutants (Fig. 2C, 2D). A longer exposure revealed that the S99P containing constructs were missing a 49 kDa band and displayed different lower molecular weight species (Fig. 2C, 2D). This is consistent with earlier data (Kedersha et al., 2016), which showed lowered expression/altered breakdown products with the original S149E mutant and a shorter piece containing the NTF2-like domain, both of which we now realize contained the additional S99P mutation.

We previously noticed that the S99P/S149E construct expressed poorly relative to the other G3BP1 variants in COS7 cells, the same cells used in the original study (Tourriere et al., 2003). To validate our U2OS findings (Fig. 1D, 1E and 2A, 2B), we then transfected COS7 cells with GFP-tagged versions of wt, S149E, S99P, and the original double mutant S99P/S149E G3BP1 (Fig. 3), and stained for the SG marker eIF3b (blue) and for sequestosome-1 (Fig. 3A-D, red), a protein adaptor that binds poly-ubiquitinylated proteins and forms cellular aggregates that promote autophagic clearance (Bjørkøy et al., 2005; Pankiv et al., 2007). SGs nucleated by G3BP1-wt or S149E are positive for eIF3b and do not contain sequestosome-1 (Fig. 3A, 3B), as expected. However, SGs nucleated by G3BP1-S99P or S99P/S149E display eIF3b-positive SGs containing subregions positive for sequestosome-1 (Fig. 3C, 3D, red arrows). To determine whether the S99P constructs were less efficiently translated, we co-transfected a neutral reporter and quantified proteins via western blotting. G3BP1 levels were normalized to reporter, and the mutants plotted relative to the normalized G3BP1-wt. The expression of the S99P variant was significantly reduced relative to WT (Fig. 3E, Fig. S3A). A short (6h) treatment with the proteasome inhibitor MG132 modestly increased its expression, but this was not significant (Fig. S3A). GFP-immunoprecipitation confirmed elevated ubiquitination in the S99P constructs (Fig. 3F). Taken together, the data indicate that S99P is unstable, ubiquinylated, and degraded (possibly in part through sequestosome-targeted autophagy), but GFP-G3BP1-S99P expression does not reduce global translation levels or transfection efficiency, as shown by the co-transfected reporter. Reduced expression may in part explain the reduced ability of G3BP1-S99P to rescue SGs (Fig. 2A, 2B) in cells lacking endogenous G3BP, but it does not explain its apparent dominant-negative effects on SGs in cells that express endogenous G3BP (Fig. 1D).

**Figure 3:**
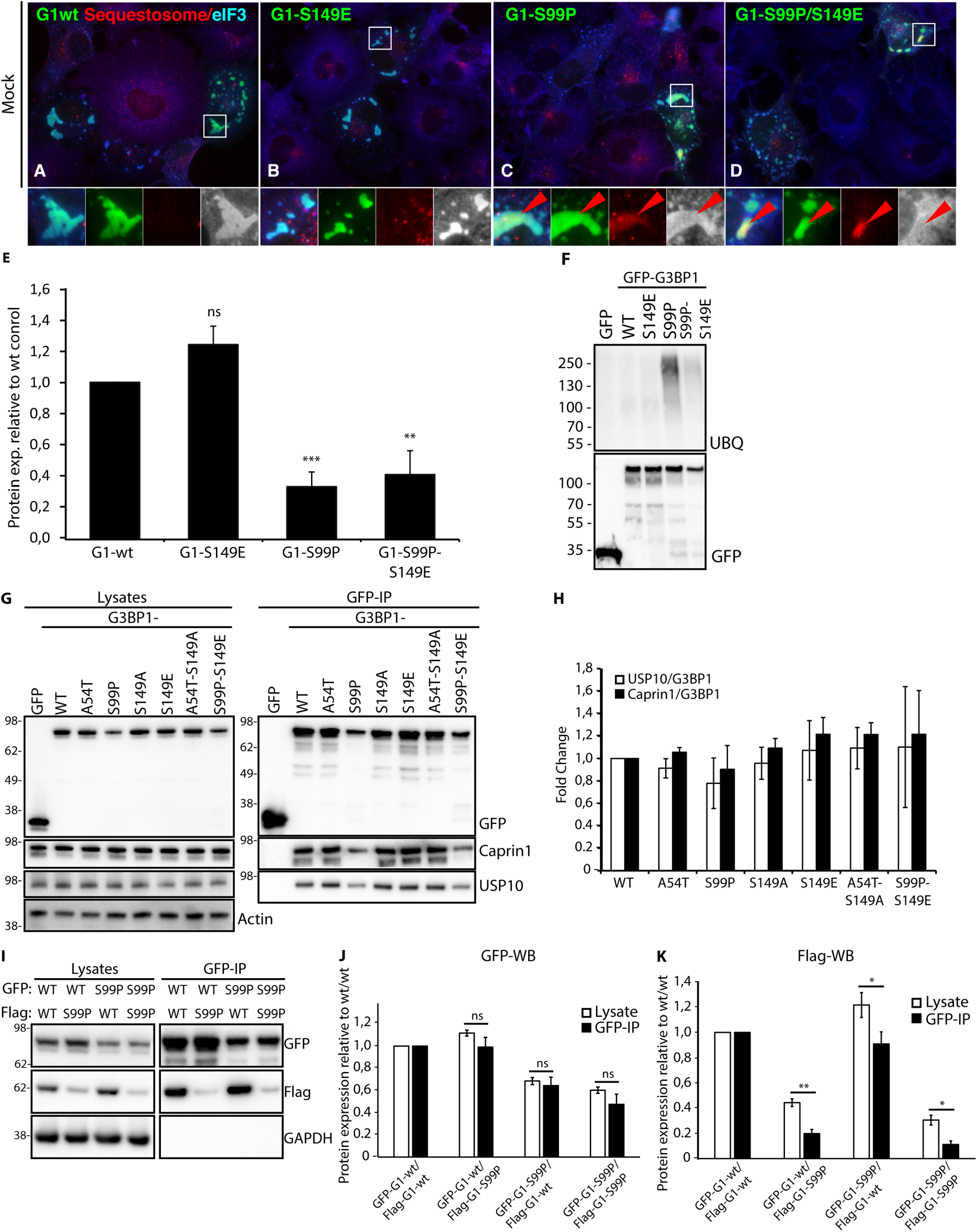
G3BP1-S99P exhibits reduced expression, increased ubiquitination, and recruits sequestosome into SGs. (A-D) COS7 cells were transiently transfected with the indicated GFP (green) constructs, fixed and stained for endogenous sequestosome/p62 (red) and eIF3b (blue). Boxed areas are shown below each panel, with colors separated (gray represents blue). Bar, 25 μm. (E) Transient co-expression of GFP-G3BP1 mutants with an HA-tagged reporter construct, quantified by western blot/densitometry for GFP and HA (Fig. S3A), expression of GFP to HA was normalized relative to the wt control, and relative values were plotted. Data shown are mean +/- SEM and analyzed using unpaired t-test. ns, not significant; **, P < 0.01; ***, P < 0.001; n = 3. (F) GFP-G3BP1 constructs transiently expressed in COS7 cells, precipitated using GFP-TRAP, and blotted as indicated. (G) Transient overexpression of indicated GFP-G3BP1 constructs in ΔΔG3BP1/2 U2OS cells, immunoprecipitated using anti-GFP and blotted as indicated. (H) Caprin1, USP10, and G3BP1 were quantified from IP Western blots (G) using densitometry and the fold change of Caprin1 or USP10 relative to GFP-G3BP1 was determined. Data shown are mean ± SEM; n = 8. (I) Transient co-overexpression of indicated GFP-G3BP1 and Flag-G3BP1 constructs in ΔΔG3BP1/2 U2OS cells, immunoprecipitated using anti-GFP and blotted as indicated. (J) GFP signal was quantified from lysates and GFP-IP Western blots using densitometry. Relative intensities to wt/wt co-overexpression were plotted. Data shown are mean +/- SEM and analyzed using unpaired t-test. ns, not significant; n = 3. (K) Flag signal was quantified from lysates and GFP-IP Western blots using densitometry. Relative intensities to wt/wt co-overexpression were plotted. Data shown are mean +/- SEM and analyzed using unpaired t-test. ns, not significant; *, P < 0.05; **, P < 0.01; n = 3.

Another property ascribed to the G3BP1-S149E phosphomimetic is increased interaction with co-expressed FLAG-Disheveled-2 (Sahoo et al., 2012); this study used the original S149E plasmid. To replicate these findings and ascertain whether they were due to the S149E or to the S99P mutation, we co-transfected our single-mutant constructs with Flag-tagged Disheveled-2 in COS7 cells, and assessed colocalization of the proteins (Fig. S3B). The results show that DVL2 only associates with G3BP1-S99P, not with G3BP1-wt or G3BP1-S149E (Fig. S3B). Remarkably, both our data and that in (Sahoo et al., 2012) show that the DVL2:G3BP1 localization is not perfectly coincident, but shows that DVL2 forms a shell around G3BP1-S99P, segregating it from the SG marker eIF3 (Fig. S3B). While the significance of this is not clear, the data suggest that the S99P mutation likely was responsible for the effects attributed to S149E.

What does the serendipitous S99P mutation tell us about G3BP-mediated SG condensation? As shown in the crystal structure (Fig. 1B) and the Ramachandran plots (Fig. S1B), the substitution of Ser-99 to a proline rigidifies the peptide backbone, which destabilizes the protein and renders G3BP1 sensitive to degradation in U2OS and COS7 cells. This could affect the ability of G3BP1 to form dimers, or to interact with proteins that normally bind to the NTF2-like domain. To distinguish between these possibilities, we first assessed effects of the S99P mutation on G3BP1 interaction with its two major partners, USP10 and Caprin1 (Kedersha et al., 2016; Panas et al., 2015; Solomon et al., 2007). GFP immunoprecipitations revealed that USP10 and Caprin1 are both co-immunoprecipitated by G3BP1 constructs containing the S99P mutation (Fig. 3G, 3H), suggesting that the USP10 and Caprin1 binding sites on G3BP are unaffected by the S99P mutation. To assess dimerization, we co-transfected ΔΔG3BP1/2 U2OS cells with various combinations of constructs expressing GFP-G3BP1-wt/S99P and Flag-G3BP1-wt/S99P. Using equal amounts of constructs, G3BP1-S99P proteins were expressed in lower levels compared to G3BP1-wt, as shown by western blot on whole cell lysates (Fig. 3I). IPs via the GFP-tag immunoprecipitated G3BP1-wt and G3BP1-S99P proteins equally well, but the lower expression of the S99P resulted in less material. As shown in Fig. 3J (quantification of 3I, left panel), the relative amounts of G3BP1-S99P to G3BP1-wt were comparable between lysates and GFP-IPs. However, proportionally less co-IPed Flag-tagged protein was detected with G3BP1-S99P present (Fig. 3K, quantification of data in Fig 3I, right panel), suggesting that G3BP1-S99P protein is impaired in interacting with G3BP1-wt protein and with itself.

### S149 is not de-phosphorylated upon arsenite treatment

Thus far, all data suggest that the S99P mutations account for the phenotype attributed to S149E, but the phosphorylation status of S149 during SG formation remains unclear. We first addressed this issue using a commercial phospho-specific antibody to the S149 site from Sigma (G8046), but found that the antibody reacted with non-phospho GFP-G3BP-S149A in G3BP1 KO cells (Fig. S3C). We then employed an IP/MS approach, in which we IP’d endogenous G3BP1 from untreated or arsenite-treated U2OS cells, then used MS to assess the phosphorylation status of two reported sites, G3BP1-S149 and G3BP1-S232 (Fig. 4A, n=4). We also examined the phosphorylation of GFP-G3BP1 stably expressed in ΔΔG3BP1/2 U2OS cells and precipitated using anti-GFP (Fig. 4B, n=6). In contrast to the original data which was obtained using CCL-39 cells (ras-transformed hamster lung fibroblasts, (Tourriere et al., 2003), we do not see a SA-induced decrease in phosphorylation at S149 in U2OS cells. On the contrary, we observe a modest but a statistically insignificant increase in S149 phosphorylation. We note that analysis of this region is particularly challenging by LCMS (Liquid Chromatography Mass Spectrometry) proteomics because of the high density of acidic residues (D and E) in the vicinity of the S149 phospho-site (YQDEVFGGFVTEPQEES#EEEVEEPEER), which confers a decreased ionization efficiency at the ElectroSpray Ionization (ESI) source, which results in poor recovery relative to other peptides analyzed by MS. To recover this peptide, we used large amounts of starting material. We attempted to assess the phosphorylation of the homologous serine in G3BP2, but failed to recover any S149 peptides, as this protein contains an even higher density of acidic residues, presumably the cause of the unreliably low recovery of the phosphopeptide by liquid chormatography/MS. We also assessed the phosphorylation of another site (S232) previously reported to be unaffected by stress (Tourriere et al., 2003), and confirm that we see no SA-induced phosphorylation changes at this site, in agreement with their findings.

**Figure 4:**
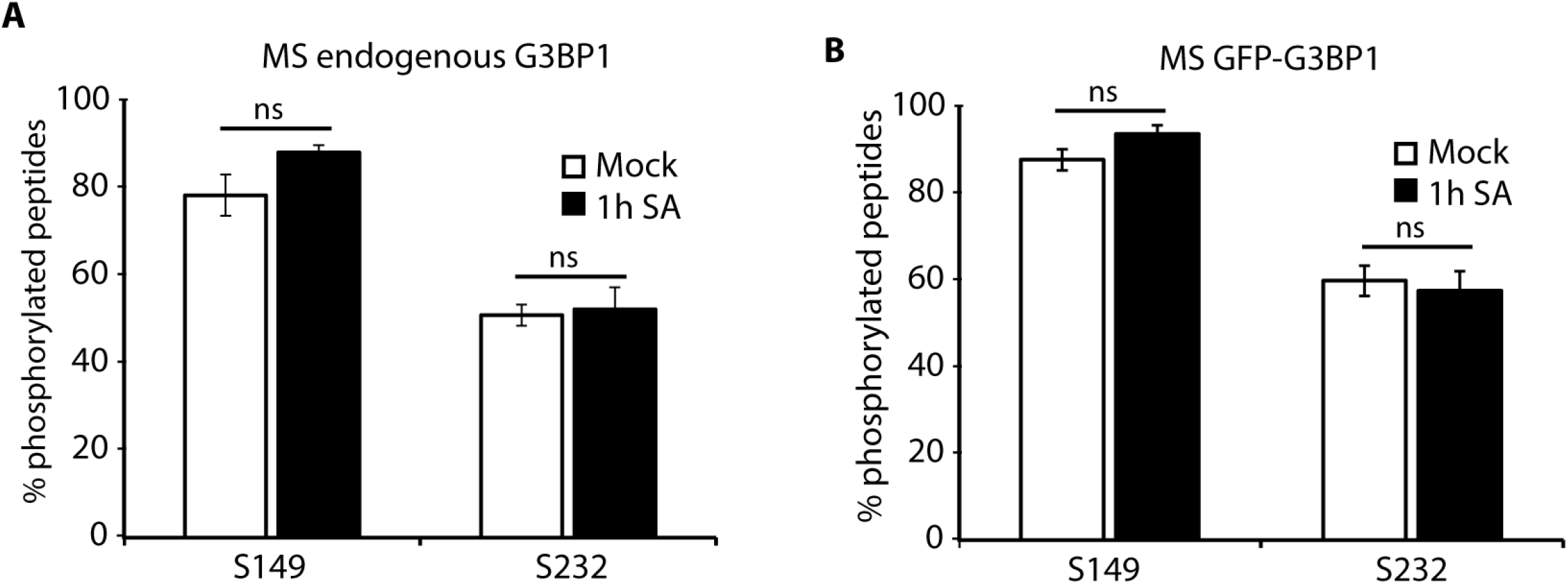
Phosphorylation of G3BP1-S149 and S232 is unaltered by SA treatment. (A) Mass spectrometry analysis of endogenous G3BP1-S149 and G3BP1-S232 phosphorylation following mock or 1h SA (500 μM) treatment. Data shown are mean ± SEM and are analyzed using the unpaired t-test. ns, not significant; n = 4. (B) Mass spectrometry analysis of stably expressed GFP-G3BP1-S149 and GFP-G3BP1-S232 phosphorylation in ΔΔG3BP1/2 U2OS cells during mock or 1h SA (500 μM) treatment. Data shown are mean ± SEM and are analyzed using the unpaired t-test. ns, not significant; n = 6.

## Discussion

G3BP is essential for SG assembly; cells lacking both G3BP1 and G3BP2 cannot assemble SGs when challenged with all known p-eIF2α dependent and some p-eIF2α independent stresses (Kedersha et al., 2016). The ability of G3BP1 to form SGs was reported to be regulated by the phosphorylation state of serine 149, such that de-phosphorylation of this amino acid promotes SG formation (Tourriere et al., 2003). In an attempt to study this posttranslational modification in more detail, we discovered additional mutations in the original constructs that were generously shared and widely distributed within the field. Specifically, we found that the pEGFP-C1-G3BP1-S149A (non-phosphorylatable) construct contains an additional alanine to threonine conversion at position 54, while the pEGFP-C1-G3BP1-S149E (phosphomimetic) construct contains an additional serine to proline mutation at position 99 (Fig. 1A). While we had originally sequenced the plasmids in 2004 to confirm the existence of the expected mutations at the S149 site, we did not sequence the entire coding region.

We created repaired constructs to contain only single mutations, then assessed their ability to nucleate SGs upon transient overexpression in U2OS cells, and for their ability to rescue SGs in ΔΔG3BP1/2 U2OS KO cells. Our data (Fig. 1, 2 and S2) show that constructs containing only S149A or S149E displayed no statistical difference in their ability to nucleate or rescue SGs, in contrast to published data (Kedersha et al., 2016; Tourriere et al., 2003). Moreover, we were unable to detect any significant SA-induced changes in the phosphorylation status of the S149 site in endogenous immunoprecipitated G3BP1 (Fig. 4A), or in GFP-G3BP1 stably expressed in ΔΔG3BP1/2 U2OS (Fig. 4B). Our data show that the S99P mutation is the sole cause of the previously described phenotype of impaired SG formation and SG rescue, and do not support the idea that phosphorylation of S149 promotes SG formation.

We mapped the exact positions of the unintentional mutations A54T and S99P in the publically available crystal structure (Schulte et al., 2016) (PDB: 5FW5) of the NTF2-like domain of G3BP1. The S99P change disrupts three hydrogen bonds within the β-sheet and a sheet-associated loop region (Fig. 1C), but also rigidifies the peptide backbone due to the structure of proline (Fig. S1B). These changes destabilize the protein to an extent that let us fail to purify the bacterially-expressed mutant protein (Fig. S1A), but more importantly render G3BP1 more sensitive to degradation when expressed in cells (Fig. 2C, 2D, Fig. 3). This is supported by *in vivo* data showing that G3BP1-S99P is less efficiently expressed (Fig. 2C, 2D, 3G, 3I), ubiquitinylated (Fig. 3F), and differently processed (Fig. 2C, 2D) relative to G3BP1-wt.

While S99P destabilizes G3BP1, it does not prevent interactions with USP10 or Caprin1 (Fig. 3G, 3H), which bind the NTF2-like domain and regulate SG formation *in vivo*. However, S99P mutations reduce dimer formation as shown by the reduced interaction of G3BP1-S99P with G3BP-wt, and the drastically reduced interaction of G3BP1-S99P with itself (Fig. 3I, 3J, 3K). As dimerization of G3BP is likely required for SG assembly, the failure of S99P mutants to interact with each other combined with their lower expression levels and ubiquitination satisfactorily explain why cells expressing G3BP1-S99P are impaired in SG assembly, and why S99P overexpression has a dominant-negative effect on SGs *in vivo*. In contrast, the single A54T mutation has only minor effects on protein expression levels, as this conserved mutation exchanges similar amino acids at a site that is located in an unstructured loop. As shown in Fig. 3G, the A54T expression levels are similar to wt-G3BP expression levels and the interactions with USP10 and Caprin1 are not impaired.

Finally, we used MS to quantify the phosphorylation status of S149 and S232 in untreated and SA treated ΔΔG3BP1/2 U2OS cells stably expressing GFP-G3BP1-wt. Previously reported findings indicating that SA treatment reduces S149 phosphorylation (Reineke et al., 2017; Tourriere et al., 2003), are not reproduced in our system. In six biological replicates, we detected no significant difference in S149 or S232 phosphorylation in response to SA treatment (Fig. 4A, 4B). Moreover, MS analysis of phophorylation of S149 and S232 of endogenous G3BP1 (Fig. 4A), and GFP-tagged stably expressed G3BP1 (Fig 4B), showed a modest but not significant increase upon SA treatment, suggesting that S149 phosphorylation does not influence G3BP’s ability to nucleate or to rescue SGs. Technical differences in the experimental setup (cell lines, antibodies) may contribute to the conflicting results, but it is important to note that the statistical significance of stress-induced reduced phosphorylation was not determined (Reineke et al., 2017; Tourriere et al., 2003).

Our findings may impact the interpretation of other studies which utilized the original pEGFP-C1-G3BP1-S149A and S149E constructs. The deacetylase HDAC6 interacts with G3BP *in vivo* and *in vitro*. Reduced binding of the S149E mutant suggested that phosphorylation modulates the HDAC6:G3BP interaction (Kwon et al., 2007), which was mapped to G3BP1 fragments containing the S149 site (the acidic region) but lacking the NTF2-like domain, hence it is possible that S149 regulates interactions with HDAC6 but not SG formation. It is notable, however, that HDAC6 has not been identified as a SG component in two major studies using proximity labeling to catalog SG components (Markmiller et al., 2018; Youn et al., 2018). Another study reported that S149E specifically interacts with Dvl2, a cytoplasmic Wnt signaling effector, but not with the non-phosphorylatable mutant S149A (Sahoo et al., 2012). Using G3BP1 fragments and co-IPs, the G3BP1:Dvl2 interaction was mapped to the NTF2-like domain and colocalization data was shown using immunofluorescence. To explain their results, the authors proposed that S149 phosphorylation changes the conformation of G3BP1 NTF2-like domain, promoting its interaction with Dvl2. While this is possible, the conformational change may be due to the S99P mutation rather than the S149E mutation, as we show that the S99P mutation mediates DVL2 colocalization in foci that lack the SG marker eIF4G1 (Fig. S3B).

Hinton et al. showed that overexpressed pseudo-phosphatase MK-STYX interacts with G3BP1 and inhibits the assembly of SGs, and first hypothesized that MK-STYX inhibition of SG assembly is G3BP-S149 dependent (Hinton et al., 2010), but subsequently showed that SG assembly is independent of S149 phosphorylation (Barr et al., 2013), using the original phosphomimetic (S149E) and non-phosphorylatable (S149A) mutants. Surprisingly, overexpression of mutationally-activated MK-STYX shows opposite effects; it induces rather than blocks SGs as does the native, enzymatically-dead version, and is capable of de-phosphorylating G3BP (Barr et al., 2013). Considering the SG-inhibitory effects of the S99P mutation in the S149E construct, it is possible that a clean S149E construct co-expressed with MK-STYX (native) would not have the same effects, as it allows the formation of SGs to the same extent as WT (Fig. 1E, 2A) or may block the formation of SGs induced by mutationally-activated MK-STYX. In the latter case, it would imply that a hypothetical blockage of SG assembly is dependent on the phosphorylation of S149. However, this is speculative and requires confirmation.

The NAD^+^ dependent deacetylase SIRT6 localizes to SGs and interacts with G3BP1, and loss of SIRT6 impairs SG formation and delays disassembly during recovery (Jedrusik-Bode et al., 2013). This study also shows that SIRT6 influences the phosphorylation of G3BP1 at serine 149, but no differences were reported on the interaction between SIRT6 and G3BP1-S149A or S149E (Jedrusik-Bode et al., 2013). This suggests that the interaction between G3BP1 and SIRT6 is independent of the NTF2-like domain and acidic region of G3BP1.

Recently, Sahoo et al., 2018 reported that in intact rat neurons translation of specific axonal mRNAs are negatively regulated by G3BP aggregation in SG like structures which can be removed by dispersion of G3BP aggregates (Sahoo et al., 2018). Using the GFP-G3BP1-S149A and S149E plasmids, the authors showed that S149E did not aggregate HuR-positive SGs, and S149E showed a higher mobility in FRAP recovery experiments. S149 phosphorylation in regenerating neurons was shown using commercial phosphospecific antibody. The authors report that S149 is de-phosphorylated upon SA treatment, again by using the S149 phosphospecific antibody, but unfortunately the data is not shown.

Rasputin, the single Drosophila ortholog of G3BP1/G3BP2, does not require phosphorylation at S142 (orthologous to mammalian S149) for SA-induced SG assembly, but phospho-S142 is required for amino-acid starvation to induce SGs (Aguilera-Gomez et al., 2017). Their data indicates that phosphorylation of rasputin/G3BP does not invariably suppress SG formation, but may actually promote it under certain conditions. Finally, casein kinase 2 (CK2) activity and S149 phosphorylation were recently linked to SG disassembly rather than assembly (Reineke et al., 2017).

It is clear that phosphorylation of endogenous G3BP occurs at S149, but the meaning of this is still unresolved. Our data indicate that S149 phosphorylation is not a simple switch that determines whether G3BP assembles SGs. It is still possible that S149 phosphorylation influences the mutually exclusive interaction of USP10 and Caprin1 in the context of recovery and disassembly of SGs, whereby the interaction of USP10 with G3BP1 is favored and USP10 can act as a “decondensase” of SGs (Kedersha et al., 2016). Alternatively, this site may influence G3BP in a signaling context, by regulating its interactions with other proteins.

## Material and Methods

### PCR mutations in pEGFP-C1-G3BP1 plasmids

The original constructs encoding pEGFP-C1-G3BP1-wt, pEGFP-C1-G3BP1-S149A and pEGFP-C1-G3BP1-S149E were a kind gift from Jamal Tazi, and were obtained from his lab on two independent occasions (July 2004, and July 2016), and sequenced throughout the coding region. Mutations were introduced into pEGFP-C1-G3BP1-wt as follows. The 5’ phosphorylated primers (see Table S1 for sequences) were mixed with 1 ng of pEGFP-C1-G3BP1-wt plasmids in a 1x Phusion PCR mastermix (Thermo Fisher Scientific) at a final volume of 25 μL. The mixture was denatured at 98°C for 30s, followed by 25 cycles of the following: 98°C for 10s, 60°C for 15s, 72°C for 2 min 30s, with a final extension step of 72°C for 5 min. 25 ng of the PCR product was ligated with T4 DNA Ligase (NEB) in a final volume of 10 μL for 1h at RT. 5 μL of the ligation mix was used for chemical transformation into high efficiency *E.coli*. Multiple clones were picked and verified by sequencing throughout the entire coding region. Sequencing reactions were carried out with an ABI3730xl DNA analyzer at the DNA Resource Core of Dana-Farber/Harvard Cancer Center (funded in part by NCI Cancer Center support grant 2P30CA006516-48).

### Bacterially-expressed protein production

The pET28-G3BP1-168-wt, S149A-(A54T) and S149E-(S99P) expression vectors were transformed into T7 express cells (NEB) and proteins were produced as described previously (Schulte et al., 2016). The poly-histidine-tagged G3BP-1-WT as well as the mutated S149A and S149E constructs were purified using immobilized metal affinity (IMAC, HisTrap FF, GE Healthcare) and the dimer was isolated using size exclusion chromatography on a Superdex 75 column (GE Healthcare).

### Structural analysis

The structure and the effect of the mutations were analyzed using PyMOL, Arpeggio, PHENIX ramalyze, Coot, as well as the Rosetta BackRuband ROSIE servers to predict the backrub models and sequence tolerance frequencies (Adams et al., 2002; Davis et al., 2006; Davis et al., 2007; Emsley and Cowtan, 2004; Lauck et al., 2010; Smith and Kortemme, 2011).

### Cell lines

COS7 cells, U2OS-wt, ΔΔG3BP1/2 U2OS, ΔΔG3BP1/2 U2OS + GFP-G3BP1-wt (Kedersha et al., 2016) cells were maintained at 5.0-7.0% CO_2_ in DMEM containing 20 mM HEPES, 10% fetal bovine serum, 100 U/mL penicillin and 100 μg/mL streptomycin.

### Transient transfection

For transient transfections, U2OS-wt and ΔΔG3BP1/2 U2OS cells (Kedersha et al., 2016) at 80-90% confluency were transfected using Lipofectamine 2000 (Invitrogen) following manufacturer’s instructions and processed after 24h. COS7 cells were transfected using SuperFect (Qiagen) and harvested after 38-42 h.

### SG induction and quantification

SGs were induced by treatment with sodium arsenite (500 μM, 1h), clotrimazole (20 μM in serum-free media, 1h), pateamine A (50 nM, 1h), or by transient transfection of SG-nucleating proteins. Cells were scored for SGs by manual counting using fluorescent microscopy, using TIA1 and eIF4G1 or eIF3b as SG markers; only cells with granules co-staining for these markers were considered SGs, and a minimum of 3 granules per cell was required to score as positive.

### Immunoblotting

Transient transfected COS7, ΔΔG3BP1/2 U2OS cells were lysed in EE-lysis buffer (50 mM HEPES, 150 mM NaCl, 2.5 mM EGTA, 5 mM EDTA, 0.5% NP40, 10% glycerol, 1 mM DTT, 0,1 mg/mL Heparin, HALT phosphatase and protease inhibitors; Thermo Fisher Scientific). Lysates were resolved in a 4-20% Mini-PROTEAN TGX Precast Gel (Bio-Rad) and transferred to nitrocellulose membranes using the Transfer-Blot Turbo transfer system (Bio-Rad). Chemiluminescent was detected using SuperSignal West Pico substrate (Thermo Fisher Scientific).

### Immunoprecipitation

100-mm dishes of 80-90%-confluent cells ΔΔG3BP1/2 U2OS cells were transiently transfected for 24h, washed with cold HBSS, and scrape-harvested at 4°C into EE-lysis buffer. Cells were rotated for 10 min at 4°C, cleared by centrifugation (10.000 g, 10 min, 4°C), and incubated with anti-GFP beads or anti-Flag-M2 affinity agarose (Sigma-Aldrich) for 2h with continuous rotation at 4°C. Anti-GFP beads were produced by expressing the GFP nanobody “Enhancer” (Kirchhofer et al., 2010) in *E.coli*, purified by size-exclusion, and coupled to cyanogen-bromide activated Sepharose (Sigma-Aldrich). Beads were washed in EE-lysis buffer and eluted directly into 2x SDS-sample buffer. Proteins were resolved in a 4-20% Mini-PROTEAN TGX Precast Gel (Bio-Rad) and transferred to nitrocellulose membranes using the Transfer-Blot Turbo transfer system (Bio-Rad) and blotted using standard procedures. Chemiluminescent was detected using SuperSignal West Pico substrate (Thermo Fisher Scientific). COS7 cells were transfected and immunoprecipitated using GFP-TRAP (ChromoTek), as described previously (Kedersha et al., 2016).

### Immunoprecipitation of GFP-G3BP1-wt for mass spectrometry

150-mm dishes of 80-90% confluent ΔΔG3BP1/2 U2OS stably expressing GFP-G3BP1-wt were left untreated or treated with 500 μM sodium arsenite for 1h. Then cells were washed with cold PBS, and scrape-harvested at 4°C into EE-lysis buffer. Cells were rotated for 10 min at 4°C, cleared by centrifugation (10,000 g, 10 min, 4°C), and incubated with anti-GFP beads for 2h with continuous rotation at 4°C. Beads were washed two times in EE-lysis buffer (500 mM NaCl) and three times in standard EE-lysis buffer (150 mM NaCl), then eluted directly into 2x SDS-sample buffer. Proteins were resolved in a 4-12% NuPAGE BT gel (Invitrogen) and stained with Coomassie Blue. GFP-G3BP1-wt bands were excised and sent for mass spectrometry. In a second experimental setup we generated G3BP1 peptides by on bead trypsin digest, skipping SDS-PAGE and Coomassie Blue staining.

### Immunoprecipitation of endogenous G3BP1 for mass spectrometry

150 mm dishes of 80-90% confluent U2OS-wt cells were treated and lysed as described above. The day before harvest respectively 150 μg of protein G beads (Pierce, Thermo Fisher Scientific) slurry was either incubated overnight with 15 μg G3BP1 (Santa Cruz, TT-Y) or 15 μg G3BP1 (Santa Cruz, H-10) or 15 μg G3BP1 (BD Transduction laboratories) in 1 mL EE-lysis buffer. Bead-immobilized G3BP1 antibody was washed once with EE-lysis buffer. Then 1 mL of U2OS-wt cell lysate was added to anti-G3BP1 (SC, TT-Y)/protein G beads and incubated for 1h at 4°C. Afterwards anti-G3BP1 (TT-Y)/protein G beads were spun down and supernatant was collected for next IP. Supernatants from the IPs were then incubated for 1h at 4°C with anti-G3BP1 (SC, H-10)/protein G beads. Then G3BP1 (H-10)/protein G beads were spun down, and supernatants collected and incubated for 1h at 4°C with anti-G3BP1 (BD)/protein G beads. All G3BP1/protein G beads were collected (450 μL total) and washed 5x with 2 mL EE-lysis buffer. Beads were then drained from washing buffer, resuspended in 500 μL in EE-lysis buffer and G3BP peptides were generated by on bead trypsin digest for mass spectrometry analysis. Lysates, combined supernatants and combined IPs were analyzed by SDS PAGE and western blotting for G3BP1 (H-10).

### SP3 based sample preparation for MS analysis

Elution from the GFP immunoprecipitation beads and endogenous G3BP1-wt IPs was done by addition of 200 μl of SDS buffer (4 % SDS, 50 mM HEPES pH 7.6, 1 mM DTT). After heating at 56°C for 5 min, the supernatant of each sample (250 μl) was transferred to a fresh tube. For cysteine blocking, 30 μl of 0.4 M chloroacetamide solution was added to each sample followed by a modified SP3 sample preparation (Hughes et al., 2014).

Briefly, two suspensions of magnetic beads (Sera-Mag Speed beads – carboxylate modified particles, P/N 65152105050250 + P/N 45152105050250, Thermo Fisher Scientific) were shaken gently until suspended. 50 μl of each suspension were combined in one tube, washed 3x with 500 μl of milliQ water and re-suspended in 500 μl of milliQ water. 50 μl of SP3 bead stock was added to each sample and gently mixed by pipetting. 400 μl of neat acetonitrile (ACN) was added to a final ACN composition of over 50%, incubated for 20 min with continuous rotation. Samples were then placed in the magnetic rack, incubated for 2 min and the supernatant discarded. The beads were washed 2x with 200 μl of 70% EtOH, with a final wash of 180 μl of neat ACN, and subsequent drying. The beads were then reconstituted in 100 μl of trypsin solution (50 mM HEPES pH 7.6, 0.5 M urea, 1 μg trypsin) and incubated for 16 h at 37°C. After incubation the supernatant was collected and transferred to a new tube, acidified with formic acid (final concentration 3%) and cleaned by reversed phase solid phase extraction (STRATA-X from Phenomenex, P/N 8B-S100-TAK) following the manufacturer’s instructions. After drying in the speedvac, samples were dissolved in 20 μl of 3% ACN, 0.1% formic acid prior to MS analysis.

### LC-MS analysis

Each sample was run in technical triplicates in addition to the biological replicates. For each LC-MS run the auto sampler (Ultimate 3000 RSLC system, Thermo Scientific Dionex) injected 5 μl of sample into a C18 guard desalting column (Acclaim pepmap 100, 75μm × 2cm, nanoViper, Thermo). After 6 min of flow at 5 μl/min, the 10-port valve switched to analysis mode in which the NC pump provided a flow of 250 nL/min through the guard column. Mobile phase A was 95% water, 5% dimethylsulfoxide (DMSO), 0.1% formic acid. The slightly concave gradient (curve 4 in Chromeleon Xpress) then proceeded from a 3% mobile phase B (90% acetonitrile, 5% DMSO, 5% water, and 0.1% formic acid) to 45% B in 65 min followed by a 10 min wash at 99% B and re-equilibration with 3% B for 5 min. Total LC-MS run time was 84 min. We used a nano EASY-Spray column (pepmap RSLC, C18, 2 μm bead size, 100 Å, 75 μm internal diameter, 50 cm long, Thermo Fisher Scientific) on the nano electrospray ionization (NSI) EASY-Spray source (Thermo Fisher Scientific) at 60°C. Online LC-MS was performed using a hybrid Q-Exactive mass spectrometer (Thermo Fisher Scientific). FTMS master scans with 70,000 resolution (and mass range 300-1900 m/z) were followed by data-dependent MS/MS (35,000 resolution) on the top 5 ions using higher energy collision dissociation (HCD) at 30% normalized collision energy. Precursors were isolated with a 2 m/z window and a 0.5 m/z offset. Automatic gain control (AGC) targets were 1e6 for MS1 and 1e5 for MS2. Maximum injection times were 100 ms for MS1 and 400 ms for MS2. The entire duty cycle lasted ~1.5 s. Dynamic exclusion was used with 30 s duration. Precursors with unassigned charge state or charge state 1 were excluded. An underfill ratio of 1% was used.

### Proteomics database search

All MS/MS spectra were searched by Sequest/Percolator under the software platform Proteome Discoverer (PD 1.4, Thermo Fisher Scientific) using a target-decoy strategy. The reference database was either the swissprot human protein database (42096 canonical and isoform protein entries, 2017-02) from uniprot.org or a database containing only the G3BP1 human protein entry. Precursor mass tolerance of 10 ppm and product mass tolerance of 0.02 Da were used. Additional settings were trypsin with one missed cleavage; Lys-Pro and Arg-Pro not considered as cleavage sites; carbamidomethylation on cysteine as fixed modification; and oxidation of methionine and phosphorylation on serine, threonine or tyrosine as variable modifications. Peptides found at 1% FDR (false discovery rate) were used by the protein grouping algorithm in PD to infer protein identities. Phosphorylation sites within a peptide sequence were assigned using the PhosphoRS node in PD. Peptide and protein areas, which serve as surrogates for the respective relative quantities across the samples, were calculated by the Precursor Ions Area Detector node in PD.

### Immunofluorescence

Cells were fixed and processed for fluorescence microscopy as described previously (Kedersha and Anderson, 2007). Briefly, cells were grown on glass coverslips, stressed as indicated, fixed using 4% paraformaldehyde in PBS for 10 min, followed by 5 min post-fixation/permeabilization in ice-cold methanol. Cells were blocked for 1h in 5% horse serum/PBS, and primary and secondary incubations performed in blocking buffer for 1h with rocking. All secondary antibodies were multi-labeling grade (tagged with Cy2, Cy3, Cy5, Jackson Immunoresearch). Following washes with PBS, cells were mounted in polyvinyl mounting media and viewed using a Nikon Eclipse E800 microscope with a 63X Plan Apo objective lens (NA 1.4) and illuminated with a mercury lamp and standard filters for DAPI (UV-2A 360/40; 420/LP), Cy2 (FITC HQ 480/40; 535/50), Cy3 (Cy 3HQ 545/30; 610/75), and Cy5 (Cy 5 HQ 620/60; 700/75). Images were captured using a SPOT RT digital camera (Diagnostics Instruments) with the manufacturer’s software, and raw TIF files were imported into Adobe Photoshop CS3. Identical adjustments in brightness and contrast were applied to all images in a given experiment.

### Drugs and chemical reagents

DMD-modified pateamine A was a kind gift from Jun Lui of Johns Hopkins. Sodium arsenite and clotrimazole were obtained from Sigma.

### Statistical analysis

Statistical analysis was performed using Microsoft Excel. Statistical differences between two groups in immunofluorescence based SG quantification, western blot experiments or mass spectrometry analysis were evaluated using unpaired Student’s t-test. p < 0.05 was considered significant. All data are expressed as mean +/- SEM.

## Supporting information

Mass Spec. dataset

## Acknowledgements

This work was supported by National Institutes of Health (GM126901) to P. Anderson, National Institutes of Health (126150) to P. Ivanov, and by the Swedish Association of Medical Research (Svenska Sällskapet för Medicinsk Forskning, SSMF) to M. Panas.

The authors declare no competing financial interests.

## Abbreviations used in this paper

MS: mass spectrometry
SA: sodium arsenite
CZ: clotrimazole
eIF: eukaryotic initiation factor
IP: immunoprecipitate
KO: knockout
Pat A: Pateamine A
p-eIF2α: phospho-eIF2α
SG: stress granule
wt: wild type
NTF2-like: nuclear transport factor 2 like
PIC: 48S preinitiation complex

---

**Table S1:**
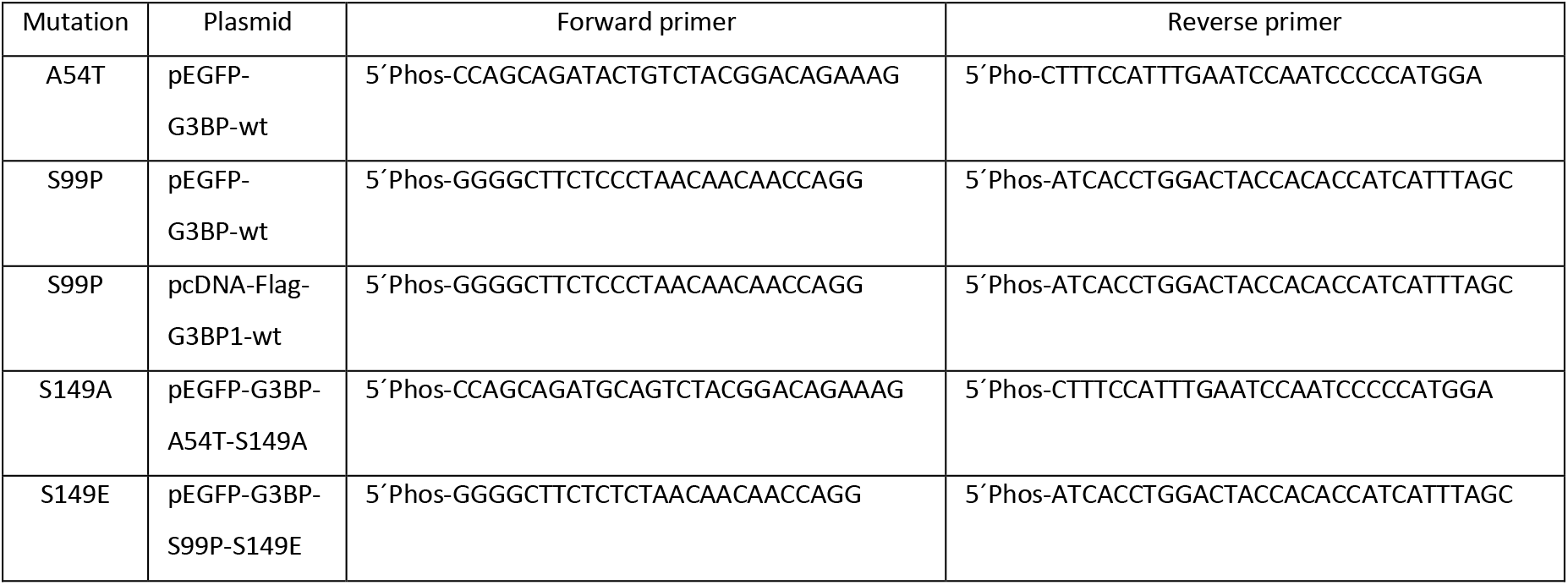
Primer sequences.

**Table S2:** List of antibodies used in this study.

| Antigen | Species | Catalog number (Clone) | Source |
| --- | --- | --- | --- |
| Actin | Mouse | Sc-8432 | Santa Cruz Biotechnology, Inc. |
| Caprin1 | Rabbit | 15112-1-AP | Proteintech Group |
| eIF4G1 | Rabbit | Sc-11373 | Santa Cruz Biotechnology, Inc. |
| eIF3b | Goat | Sc-16377 (N-20) | Santa Cruz Biotechnology, Inc. |
| Flag | Mouse | F3165 | Sigma-Aldrich |
| GAPDH | Mouse | Sc-47724 | Santa Cruz Biotechnology, Inc. |
| G3BP1<br>Epitope | Mouse | Sc-81940 (TT-Y)<br>Recombinant hG3BP1 | Santa Cruz Biotechnology, Inc. |
| G3BP1<br>Epitope | Mouse | Sc-365338 (H-10)<br>aa 158-251 hG3BP1 | Santa Cruz Biotechnology, Inc. |
| G3BP1<br>Epitope | Mouse | 611126<br>aa 210-323 hG3BP1 | BD Transduction Laboratories |
| phos-G3BP1-S149 | Rabbit | G8046 | Sigma-Aldrich |
| GFP | Rabbit | ab290 | Abcam |
| GFP | Chicken | G160 | ABM |
| HA | Mouse | MMS-101R | Covance |
| Sequestosome 1 | Mouse | Sc-28359 | Santa Cruz Biotechnology, Inc. |
| TIA1 | Goat | sc-1751 | Santa Cruz Biotechnology, Inc. |
| USP10 | Rabbit | A300-900A | Bethyl Laboratories |
| USP10 | Rabbit | A300-901A1 | Bethyl Laboratories |
| Ubiquitin | Mouse | Sc-271289 | Santa Cruz Biotechnology, Inc. |

**Figure S1:**
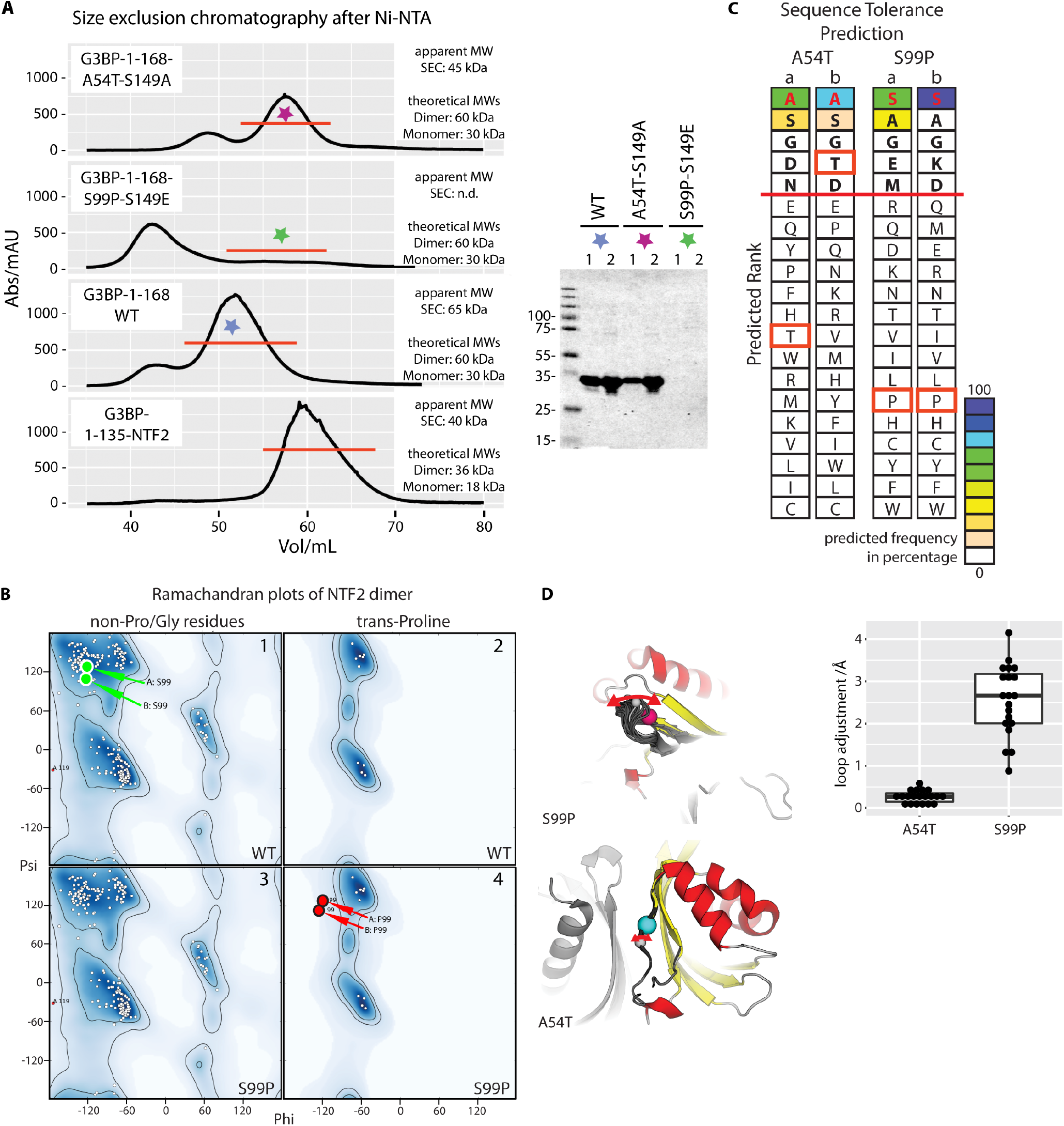
(A) Size exclusion and SDS-PAGE analysis of heterologously produced G3BP1-1-168 wild-type, S149A (unintentional A54T) and S149E (unintentional S99P) constructs. The SEC profile of the NTF2-like G3BP1 domain (sequence-derived MW is 18 kDa) used for crystallization is shown for comparison. Wild-type G3BP1-1-168 (30 kDa) yielded a major peak at a retention volume of about 52 mL, corresponding to an apparent MW of 65 kDa, in agreement with the theoretical size of the dimer. The S149A (A54T) mutant eluted at about 58 mL corresponding to an apparent MW of 45 kDa, in-between the monomer and dimer. For the S149E (S99P) construct a major peak at the void volume of the column was obtained, indicating aggregated protein. SDS-PAGE analysis showed protein bands at the expected MW of about 30 kDa for the combined major peak fractions of G3BP1-wt and S149A (A54T). We did not detect any S149E (S99P) protein in the indicated fractions that were chosen based on the purification of the wild-type and S149A (A54T) protein. The major “void” peak was not analyzed in SDS-PAGE. Numbers 1 and 2 indicate before and after concentrating the combined fractions using ultracentrifugation. (B) *In silico* substitution of Ser-99 to Pro in monomer A and B shifts the formerly favored Phi/Psi angles of Ser (top 1, green arrows) into disallowed regions in the proline Ramachandran plot (below 4, red arrows). (C) The ranked table of residues with their respective predicted frequencies illustrates that both wild-type residues at positions 54 and 99 are favored in the two chains of the NTF2 dimer (a, b). The red line indicates a typical cutoff of picking the top 5 amino acid choices at each position. While threonine (red boxes) appears among the five highest ranked residues in one chain of the dimer, proline at position 99 (red boxes) is listed at the lower end of the ranked table of both chains. (D) ‘Backrub’-modeled structures of S99P mutants reveal a 2.5 Å perturbation of the Ser-99 associated loop. The models of A54T are almost identical to the wild-type structure with perturbations in the range of 0.2 Å. Cartoons were drawn and color-coded as in Figure 1B. The perturbation of the loops was measured as point-to-point distances of the highlighted wild-type Cα-atom (grey sphere) to the respective atom in the mutant structures.

**Figure S2:**
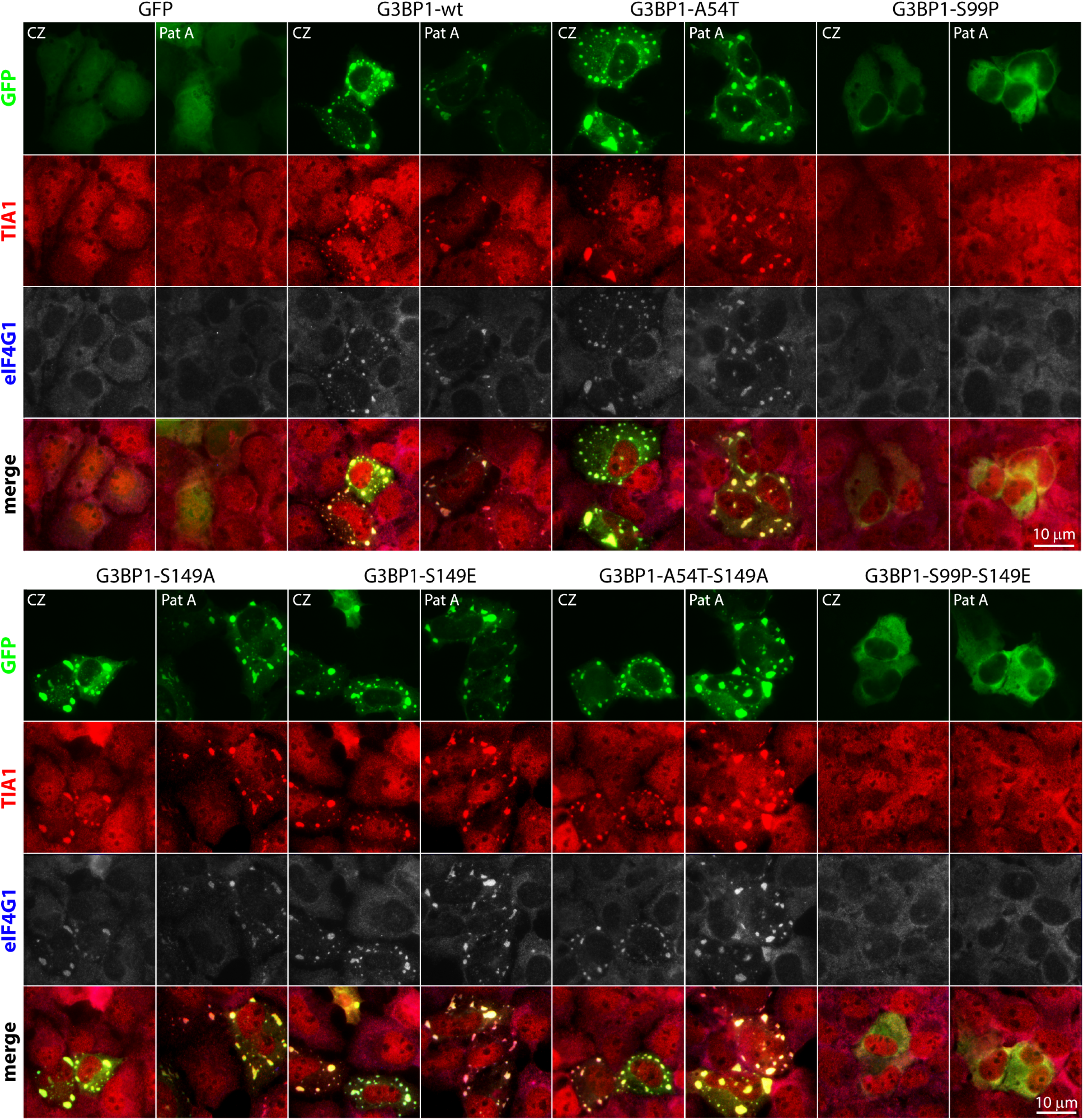
GFP-G3BP1-S99P and GFP-G3BP1-S99P-S149E mutants display reduced SG rescue. ΔΔG3BP1/2 U2OS cells transiently transfected with the indicated GFP (green) constructs, then treated with 20 μM clotrimazole (CZ) or 50 nM pateamine A (Pat A) for 1h. Cells were fixed and stained for endogenous TIA1 (red) and eIF4G1 (blue represented as grey). Bar, 10 μm.

**Figure S3:**
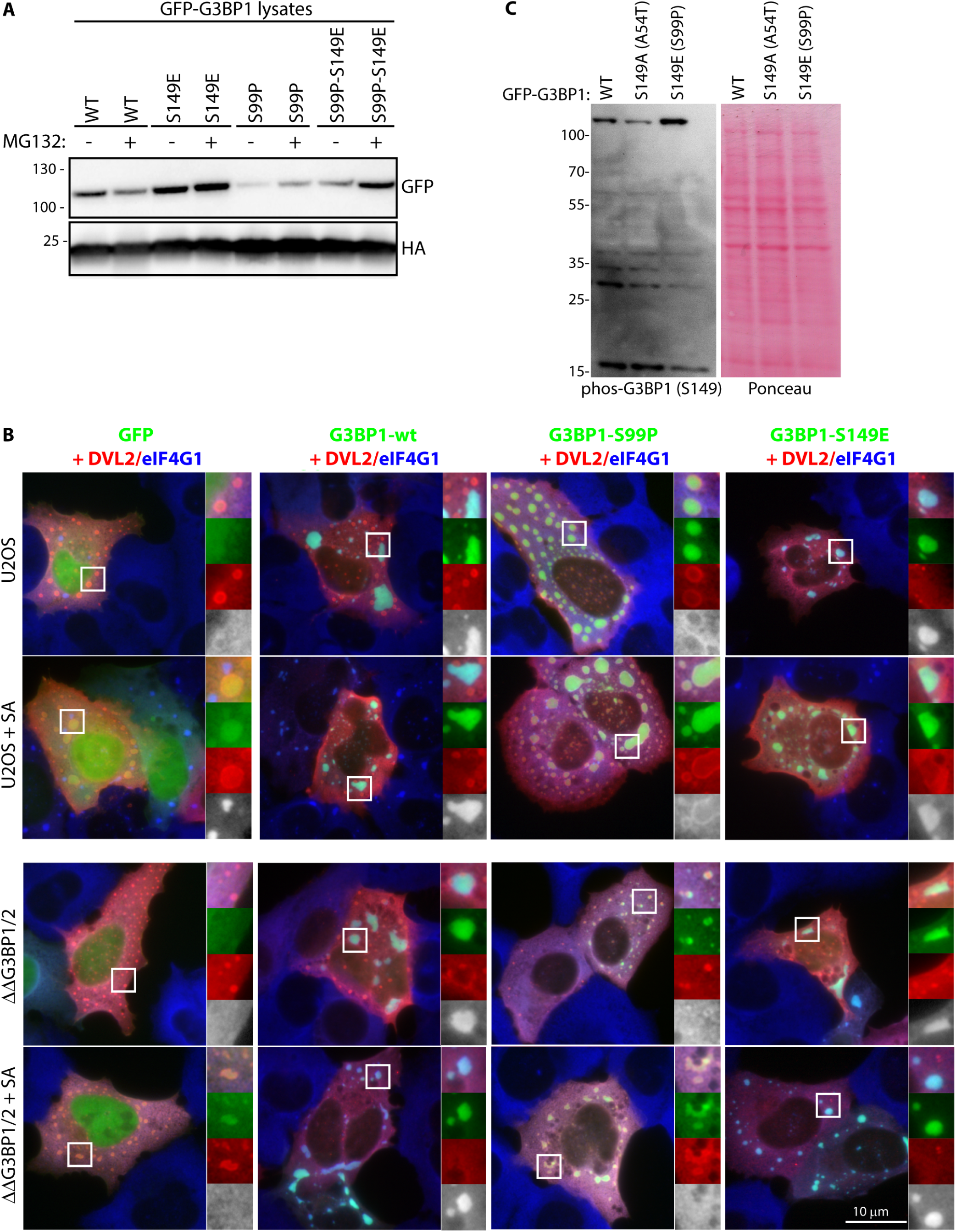
Overexpressed Dvl2 forms shells around G3BP1-S99P and does not recruit eIF4G1 in ΔΔG3BP1/2 U2OS cells. (A) Transient co-expression of GFP-G3BP1 mutants with an HA-tagged reporter construct, untreated or treated with MG132 for 8h, blotted as indicated. (B) GFP-tagged forms of G3BP1 (green) were co-transfected with Flag-Dvl2 (red) in U2OS (top 2 rows) or ΔΔG3BP1/2 U2OS (bottom 2 rows) unstressed or stressed with SA (500 μM) for 1h. Cells were counterstained for eIF4G1 (blue). Bar, 10 μm. (C) ΔG3BP1 U2OS cells stably expressing GFP-G3BP1-wt, S149A (A54T), S149E (S99P). Lysed and separated by SDS PAGE, membrane stained with ponceau (right panel) and then blotted for phos-G3BP1-S149 (Sigma, G8046, left panel).

